# An integrated, modular approach to data science education in the life sciences

**DOI:** 10.1101/2020.07.25.218453

**Authors:** Kimberly A Dill-McFarland, Stephan G König, Florent Mazel, David Oliver, Lisa M McEwen, Kris Y Hong, Steven J Hallam

## Abstract

We live in an increasingly data-driven world, where high-throughput sequencing and mass spectrometry platforms are transforming biology into an information science. This has shifted major challenges in biological research from data generation and processing to interpretation and knowledge translation. However, post-secondary training in bioinformatics, or more generally data science for life scientists, lags behind current demand. In particular, development of accessible, undergraduate data science curricula has potential to improve research and learning outcomes and better prepare students in the life sciences to thrive in public and private sector careers. Here, we describe the Experiential Data science for Undergraduate Cross-Disciplinary Education (EDUCE) initiative, which aims to progressively build data science competency across several years of integrated practice. Through EDUCE, students complete data science modules integrated into required and elective courses augmented with coordinated co-curricular activities. The EDUCE initiative draws on a community of practice consisting of teaching assistants, postdocs, instructors and research faculty from multiple disciplines to overcome several reported barriers to data science for life scientists, including instructor capacity, student prior knowledge, and relevance to discipline-specific problems. Preliminary survey results indicate that even a single module improves student self-reported interest and/or experience in bioinformatics and computer science. Thus, EDUCE provides a flexible and extensible active learning framework for integration of data science curriculum into undergraduate courses and programs across the life sciences.

**Availability and implementation:** The EDUCE teaching and learning framework is accessible at educe-ubc.github.io

## Introduction

We live in an increasingly data-driven world, where high-throughput sequencing and mass spectrometry platforms have generated a veritable tsunami of multi-omic information (*e.g*. DNA, RNA, protein, and metabolite) spanning multiple levels of biological organization [1, 2]. For example, over 31 terabases of genomic sequence information are created on average per second, and rates are expected to continue to increase with continued technological improvement across the life sciences [3]. In this light, major challenges in the life sciences are shifting away from data generation and processing to interpretation and knowledge translation resulting in a need for increased training in bioinformatics or more generally, data science.

Despite calls to action as early as 20 years ago [4], there remains a sustained, unmet need for bioinformatics training in the life sciences [5]. In 2015, Horton and Hardin described prevalent challenges and opportunities of this mandate in a special issue of the American Statistician focused on teaching statistics students how to “Think with data” [6]. Beyond reforming curriculum in mathematics and statistics, a meta-analysis of surveys from organizations such as the Global Organisation for Bioinformatics Learning, Education and Training (GOBLET), the European life-sciences Infrastructure for Biological Information (ELIXIR), the U.S. National Science Foundation (NSF), and the Australia Bioinformatics Resource (EMBL-ABR) found the desire for training to be widespread both in terms of geography and career level [5]. While a number of training formats were applicable, the most common ask for undergraduate students was integrated data science training within current degree programs [5]. Such an integrated program would ensure that all students develop core competency needed for continued studies and career development.

In order to meet this clear and present need for data science education in the life sciences, the University of British Columbia (UBC) has launched a number of initiatives in recent years. These include the Master of Data Science program as well as several specialized undergraduate majors like biotechnology (BIOT) and combined computer science-life science programs. While these programs provide a subset of students with in-depth training, they are inherently self-selecting and limit integrated and inclusive development of core competency across the faculty of science [7]. Given that data science training is necessary for life science graduates and specialized degree programs do not attract all students, we posit that such training needs to be incorporated into the fabric of undergraduate coursework such that students are able to progressively develop confidence and skills in an active learning process [8, 9].

With this teaching and learning paradigm in mind, the Experiential Data science for Undergraduate Cross-Disciplinary Education (EDUCE) initiative was launched in Fall 2017. The following provides an in-depth description of the EDUCE framework as well as preliminary findings from EDUCE modules implemented in Microbiology and Immunology (MICB) courses at UBC.

## Experiential Data science for Undergraduate Cross-Disciplinary Education (EDUCE)

EDUCE seeks to develop an extensible, progressive, cross-disciplinary and collaborative framework to equip undergraduate students in the life sciences with core competency in data science. EDUCE curriculum provides students with the most commonly needed data science skills as defined by recent surveys [5]. Focal learning objectives of the EDUCE teaching and learning framework ask students to learn to:

- Recognize and define uses of data science
- Explore and manipulate data
- Visualize data in tables and figures
- Apply and interpret statistical tests

Sub-objectives are defined using discipline-specific data sets, questions, and software. The combined set of learning objectives are achieved through 1) a modular curriculum plugged into existing courses, 2) coordinated co-curricular activities, and 3) a cross-disciplinary community of practice that brings together teaching teams across multiple training levels. The resulting teaching and learning framework allows students to progressively develop core competency over several years of integrated practice.

### Modular curriculum

The EDUCE framework is modular, providing for flexible integration of data science curriculum within a single course or a series of interconnected courses. Modules are self-contained, adjustable bundles of learning materials that can be delivered as stand alone classes, recurring instances e.g. ‘data science Fridays’, or integrated consecutively in support of capstone projects.

Because EDUCE is modular, it overcomes several of the largest barriers to data science education in the life sciences including instructor capacity, student prior knowledge, and relevance to discipline-specific problems. [10]. Specifically, modules are developed and delivered using a community of practice model that empowers instruction across different training levels and disciplines [11]. Teaching assistants (TAs), postdocs, instructors and research faculty co-create learning materials and work together across courses to implement EDUCE modules resulting in a zone of proximal development for curriculum development [12]. EDUCE modules assume no prior knowledge as they incorporate all necessary introductory and background material, including materials unique to that module and review activities from previous modules. Finally, module integration circumvents traditional course creation, *i.e*. new course codes, thus allowing quicker, more broadly accessible curriculum deployment within the university ecosystem.

In addition, modular instruction allows for more timely and direct linkages to discipline-specific content than traditional course structures. For example, students cover the global impacts, thermodynamics, etc. of the nitrogen cycle in their regular course content and then immediately enter an EDUCE module in which they quantify and plot nitrogen species in the ocean using R [13]. This aids student learning in both the discipline and data science as students are able to reference prior knowledge [14] while reflecting on the meaning and application of a nascent skill [15]. Thus, modules leverage student interest in their chosen area of study and maximize the relevancy of data science content used in the learning process.

An EDUCE module consists of all materials related to a set of learning objectives and includes 1) introduction, 2) practice, 3) application, and 4) communication (Fig 1). As students progress from introductory to advanced modules across several courses, the teaching and learning focus transitions from introduction and practice to application and communication. Thus, while each module contains all four aspects, later modules build on prior knowledge to both reinforce key concepts and stimulate students to apply developing data science skills to higher dimensional problems with emphasis on visualization, interpretation and communication. This builds progressive competency in data science over several years of instruction without the need for additional course requirements.

**Fig 1.**
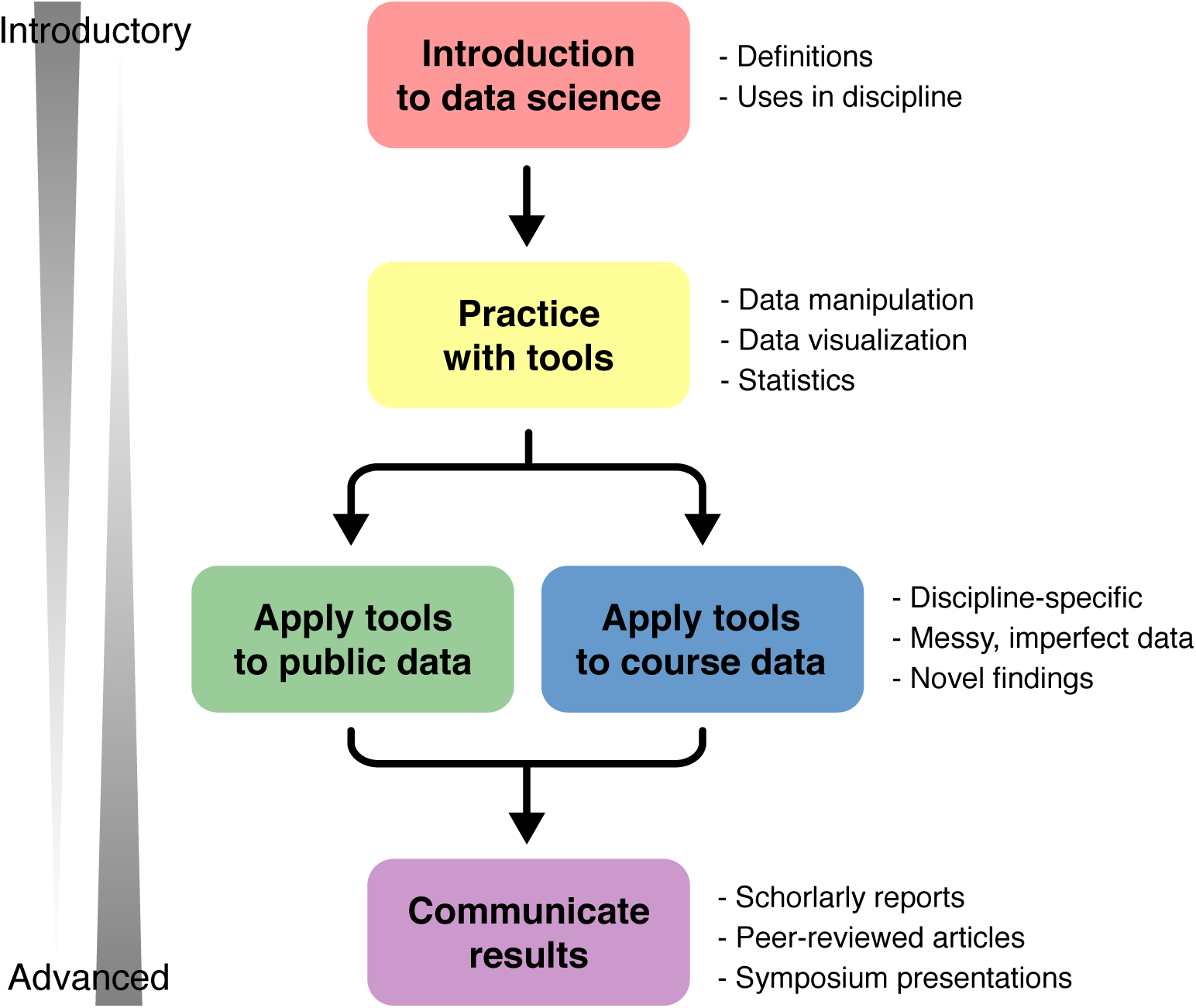
EDUCE module overview. Each EDUCE module consists of introduction, practice, application, and communication. Early, introductory modules are focused on introduction and practice while later, more advanced modules challenge students to apply their developing data science skills to higher dimensional data with emphasis on visualization, interpretation and communication.

Modules vary in size from a single activity to capstone projects, and they can include a range of materials bundled together based on learning objectives, available class time, and instructor needs. Examples of specific MICB modules are shown in detail below and additional examples can be found in the “Course Compliler” app on the EDUCE organizational website (educe-ubc.github.io/compiler.html).

- Lecture slides
- Notes / handouts
- In-class tutorials
- At-home tutorials
- Screen capture videos
- Individual assignments
- Team projects

### Example modules in Microbiology

EDUCE modules have been integrated into seven third and fourth year MICB courses at UBC selected based on internal curriculum review and instructor interest (Table 1). These courses include lectures, wet labs, and dry labs (*i.e*. tutorials).

**Table 1.** EDUCE courses in microbiology (MICB)

| MICB code | Name | L | W | D | Hrs/wk | EDUCE hrs |
| --- | --- | --- | --- | --- | --- | --- |
| 301 | Microbial ecophysiology | X |  |  | 3 | 5 |
| 322 | Molecular microbiology laboratory | X | X |  | 6 | 3 |
| 323 | Molecular immunology and virology laboratory | X | X |  | 7 | 3 |
| 405 | Bioinformatics | X |  | X | 4 | 17 |
| 425 | Microbial ecological genomics | X |  |  | 3 | 17 |
| 421, 447 | Experimental microbiology/molecular biology (CURE) | X | X |  | 7 | 2+ |

L: Lecture, W: Wet lab, D: Dry lab, CURE: Course-based undergraduate research experience

EDUCE content varies in each course and builds from the third (300+) to fourth-year (400+) (Fig 2). Ideally, students enter the program in MICB 301 where they master basic R/RStudio scripting to manipulate data, create figures, and perform *t*-tests. Then, students acquire additional practice in R/RStudio in 300-level labs (MICB 322 and 323) where they employ specialized R packages to manipulate and plot data generated in the course. For example, MICB 322 uses the Sushi package [16] to visualize transposon insertions in the *Caulobacter* mutant library created by students in the course. Next, students progress to elective 400-level courses including MICB 405 and 425, and course-based undergraduate research experiences (CUREs) including MICB 421 and 447. In the elective courses, EDUCE modules are built into the curriculum and include capstone projects using published but under-explored metagenomic and metatranscriptomic data sets [17]. In contrast, CUREs involve student-led research projects that generate novel data. Thus, EDUCE serves as a consultation resource to help students with R/RStudio packages, Unix tools, or statistical methods as required for their individual projects. All EDUCE elective courses offer the opportunity for students to communicate their scientific findings through publication in the Undergraduate Journal of Microbiology and Immunology (UJEMI, jemi.microbiology.ubc.ca) and/or presentation at the MICB Undergraduate Research Symposium (URS, jemi.microbiology.ubc.ca/UndergraduateResearchSymposium).

**Fig 2.**
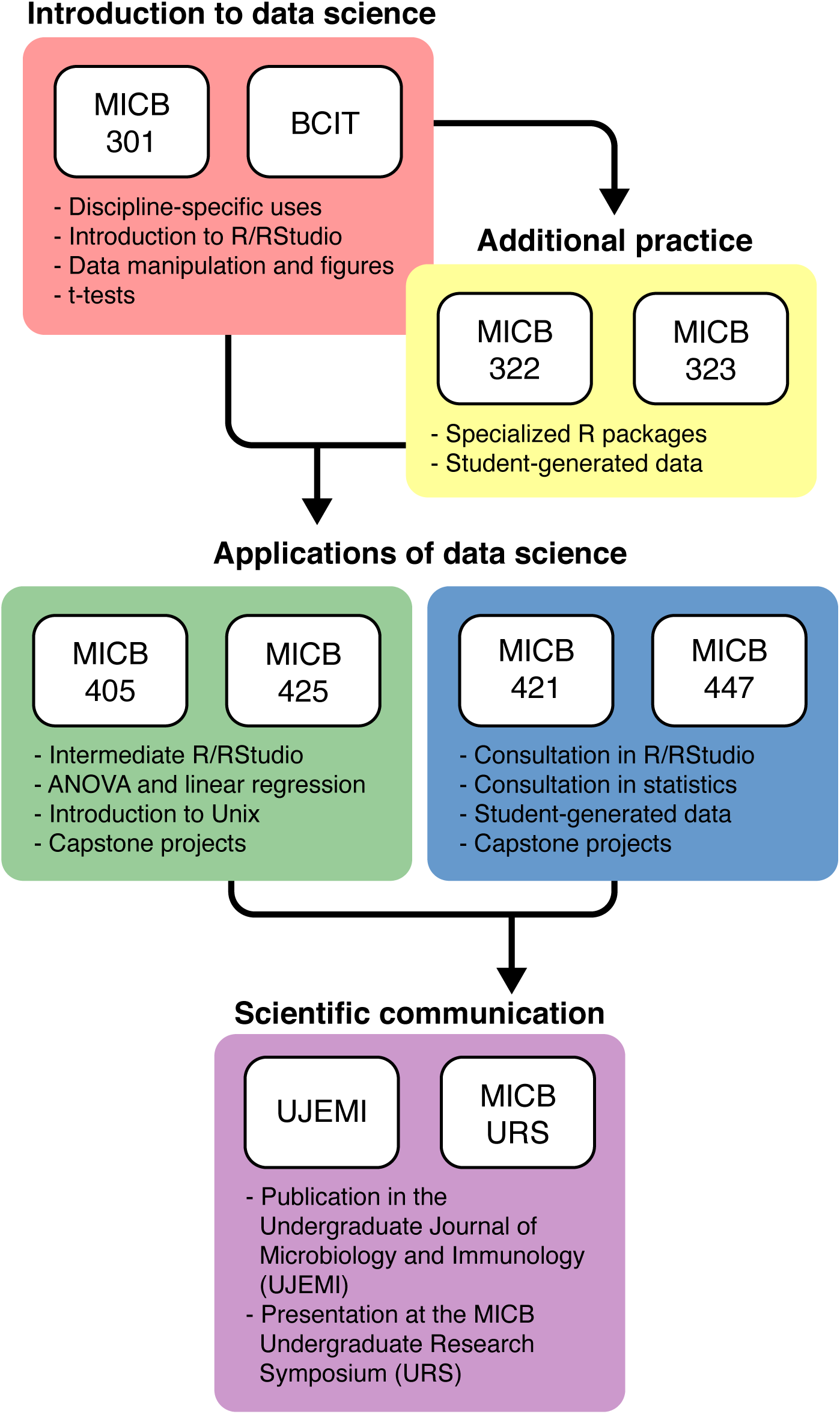
EDUCE curriculum in microbiology. Students enter the program in MICB 301 or equivalent coursework at the British Columbia Institute of Technology (BCIT). Some students take additional 300-level courses, MICB 322 and 323. Then, all students progress to one or more elective 400-level courses with prescribed outlets for communication of results. MICB: microbiology

While the above is an ideal module progression, the EDUCE framework is intentionally adjustable. This is necessary, because MICB prerequisite structure and major requirements do not resolve a linear progression for all students. Microbiology Honours (MBIM) and MICB degrees require the majority of EDUCE courses, while combined majors require a subset of EDUCE courses with little overlap between elective options. Specifically, the MICB/MBIM and Computer Science (+CPSC) or Earth, Ocean, and Atmospheric Sciences (+EOAS) majors require the first EDUCE course (MICB 301) but different 400-level elective courses (MICB 405 and 425, respectively). MICB/MBIM + EOAS also requires an additional 300-level EDUCE ‘practice’ course. The Biotechnology (+BIOT) degree, on the other hand, is a joint program wherein students spend their second and third years at BCIT. Thus, +BIOT does not require any 300-level EDUCE courses. In addition, many students choose to take additional 400-level elective courses beyond those required by their major (Table 2).

**Table 2.** EDUCE course requirements for microbiology majors.

| MICB code | Undergraduate major |  |  |  |
| --- | --- | --- | --- | --- |
|  | MICB/MBIM | +CPSC | +EOAS | +BIOT |
| 301 | R | R | R | S |
| 322 | R | S | R | O |
| 323 | R | S | O | O |
| 405 | O | R | S | R |
| 425 | O | O | R | S |
| 421/447 | R | O | O | S |

R: Required, S: Suggested, O: Optional. MICB: Microbiology, MBIM: Microbiology Honours, CPCS: Computer Science, EOAS: Earth, Ocean, and Atmospheric Sciences, BIOT: Biotechnology.

These course modules incorporate the four focal learning objectives and form the basis of the EDUCE teaching and learning framework. While all MICB students obtain some level of data science competency through EDUCE modules within required courses, not all students are exposed equally due to course dependency relationships or specialized program requirements. To help overcome this variation, a persistent co-curricular layer was developed that is coordinated in time with EDUCE module deployment.

### Co-curricular activities

EDUCE co-curricular activities reinforce and expand on course modules by providing undergraduate students with multiple data science learning opportunities outside of the classroom setting. Similar to course modules, co-curricular activities are flexible and integrated to provide students with progressive levels of instruction from introductory to advanced (Fig 1). Such activities include workshops, hackathons, and directed studies that incorporate one or more focal learning objectives and meet the following criteria.

They must first and foremost be accessible in terms of cost, content, and schedule. Specifically, undergraduate students are often unable to pay the high costs of data “boot camps” [18] and resist outside training opportunities due to motivational barriers or limited time [19]. Secondarily, EDUCE co-curricular activities directly reiterate and build on learning objectives from related course modules. This feature is intended to improve student learning by referencing prior knowledge [14] and imparting discipline-specific meaning to developing skills [15]. This is particularly important for short-term training opportunities as recent research suggests that fully stand-alone programs, such as “boot camps”, do not promote long-term skill development even at the graduate level [20, 21]. Finally, co-curricular activities invoke the same or similar data sets and packages as course modules.

Course modules are coordinated in time with co-curricular activities to prime student participation. For example, in the Fall term introductory R workshops are offered at the same time that students are introduced to R scripting in MICB301 and provide a refresher for students in MICB405 prior to learning more advanced functions and faceted data visualization. While integration with classroom module requires that co-curricular activities are discipline-specific this does not necessarily exclude other participants. For example, an R [13] workshop using oceanic oxygen data may be designed for life or Earth science students but is no less accessible to other disciplines than the commonly used R “cars” data [22]. While current EDUCE workshops focus on applying data science skills in ecology and microbiology, thematic workshop can be readily developed to focus on discipline-specific data sets or software applications.

EDUCE co-curricular activities fulfill a number of roles. Workshops provide a relaxed and open environment in which to practice current course modules or review previous ones and allow students opportunities to explore learning modules from courses that conflict with their registered schedules. Workshops also provide the teaching team an opportunity to benchmark new curriculum prior to module integration into classroom settings. Other co-curricular activities such as hackathons provide opportunities for students to work in a more social team setting to solve interesting problems with real world implications while directed studies provide students with the opportunity to use their data science skills to answer specific research questions. During workshops and hackathons, substantial time is dedicated to promoting scaffolding between participants across different training levels.

### Example co-curricular workshops in Microbiology

The EDUCE program at UBC has partnered with the Ecosystem Services, Commercialization Platforms and Entrepreneurship (ECOSCOPE, ecoscope.ubc.ca) training program and the Applied Statistics and Data Science Group (ASDa, asda.stat.ubc.ca) to deliver data science workshops accessible to participants across different training levels from undergraduate students to industry professionals. The current workshop portfolio (github.com/EDUCE-UBC/workshops_access) consists of 32 hours of content in R [13] including the same oceanic geochemical data set [23] used in most EDUCE course modules. These workshops incorporate the four focal learning objectives and include:

- Introduction to R
- The R tidyverse
- Reproducible research in R and Git
- Intermediate R programming
- Statistical models in R
- Phylogenetics and microbiomes in R

Introduction to R is a short refresher of the MICB 301 module or an introduction to the workshop series for participants not enrolled in EDUCE courses. The R tidyverse closely follows its respective module in MICB 405 and 425 (Fig 1) and thus, can be used as a refresher or additional practice. The remaining workshops have some overlap with course modules (like ANOVA in Statistical models in R and MICB 405) but mainly build on the fourth year modules to challenge students to continue developing their data science skills. In 2020, two advanced workshops were added including *Introduction to programming and plotting in Python* and *Visualization of metagenomic and metatranscriptomic data*.

Importantly, ECOSCOPE sponsorship allows undergraduates to take any workshop for 10 CAD as compared to regular registration fees of 100 CAD or more. This, along with careful scheduling to 1) align workshop content with undergraduate course modules and to 2) accommodate the most common undergraduate schedules in MICB, facilitates increased undergraduate participation. Workshops are actively advertised in undergraduate MICB courses, the ECOSCOPE website and multiple campus list serves. If scheduling conflicts prevent a significant number of undergraduates from participating, additional workshop sessions are planned on an ad hoc basis. While current workshops run on a subsidized cost recovery basis this model can only be maintained with sustained institutional support that ideally would enable registered students to participate for free.

### Community of practice

Data science is an inherently cross-disciplinary field traditionally comprised of statistics, mathematics, and computer science. EDUCE builds on this tradition through a cross-disciplinary community of practice including traditional and less conventional fields. While EDUCE is coordinated by a research faculty member and postdoctoral teaching and learning fellow (TLF) appointed through MICB, the teaching team spans 10 departments across three faculties (Fig 3). For example, statistics consults on module development, mathematics provides resource support, and teaching assistants (TAs) have been recruited from the Faculties of Science, Applied Science, and Medicine. The community consists of multiple training levels including undergraduate and graduate students, postdoctoral fellows, instructors and research faculty. Overall, the EDUCE teaching team brings together expertise from across the university to provide relevant curriculum grounded in both current data science best practices and scholarship of education leadership.

**Fig 3.**
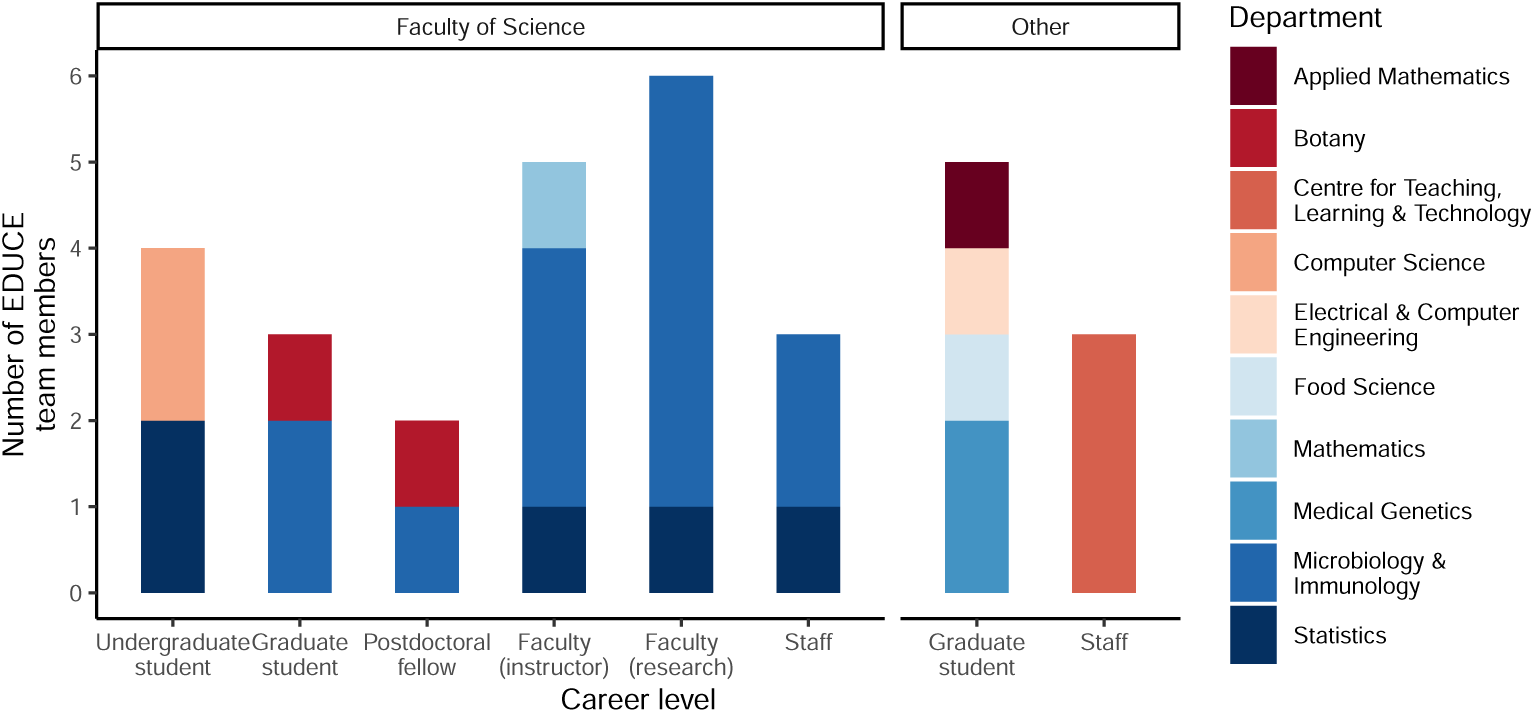
EDUCE team members at UBC. The EDUCE initiative is led by MICB in the Faculty of Science. However, TAs, consultants, collaborators, and other support come from across the campus community and include 10 departments from 3 Faculties (Science, Applied Science and Medicine). Team members include multiple training levels that work together to develop and deploy content in both courses and co-curricular activities.

The EDUCE community of practice reflects widespread changes in teaching in MICB. A growing number of MICB courses (24% in 2018/19) are taught by a team of faculty often including both tenure-track researchers and instructors. This team teaching brings additional expertise, styles, and perspectives into the classroom [24, 25] and encourages collaboration and community [25]. In addition to faculty, EDUCE TAs contribute to team teaching. Unlike traditional appointments, EDUCE TAs are not assigned to a specific course nor do they always come from the same department. Instead, EDUCE TAs are recruited from across the campus community and participate at every level, from curriculum development to office hours and instruction. This provides undergraduate students in EDUCE courses with diverse resources, viewpoints, and role models as they progress through modules and co-curricular activities. At the same time, EDUCE TAs gain hands-on teaching experience and evaluation to build their individual teaching portfolios.

## Preliminary outcomes of EDUCE

EDUCE launched at UBC in Fall Term 1 of the 2017/18 academic year. At present, approximately 475 (redundant) students are exposed to EDUCE modules through 45 total classroom hours in five MICB courses per year. Over the first two years, co-curricular workshops were taken by 77 EDUCE students, which resulted in increasing undergraduate participation from 0% (2016/17, prior to partnership with EDUCE) to 8.8% (2017/18) to 23% (2018/19) of total registrants. Undergraduate participants in workshops appears to be stabilizing around 20% in 2019/20.

Evaluation of EDUCE is on-going and includes student surveys with Research Ethics Board Approval (H17-02416) at the start (pre) and end (post) of courses (Text S1) as well as at the end of workshops. Courses with <5 EDUCE hours (Table 1) were excluded to avoid redundant surveying of students. For example, >95% of students in MICB 322 concurrently take MICB 301. At present, 439 pre-course responses and 400 post-course responses have been collected with 91 students indicating willingness to complete post-graduation surveys or focus groups. An additional 44 responses have been collected from co-curricular workshops.

Preliminary analysis of the first EDUCE course, MICB 301, from 2017-18 revealed that student self-reported interest in ‘bioinformatics’ increased with a significant transition from medium to high interest (Monte Carlo FDR P = 1.82E-2, Fig 4). Similarly, student self-reported experience in ‘bioinformatics’ significantly increased from none to low (FDR P = 1.84E-7) or medium (FDR P = 2.74E-5) as well as from low to medium (FDR P = 2.69E-2). This is relevant to attracting more students to fourth year elective courses implementing EDUCE modules including MICB405 and MICB425. In contrast, self-reported interest in ‘computer science’ remained constant but showed significant gains in experience from none to low (FDR P = 2.44E-3), and ‘statistics’ had no significant changes in either interest or experience. Thus, it appears that even minimal exposure to data science in disciplinary coursework (5 hours, (Table 1)) may impact student interest and confidence in some data science areas. More importantly, at least one case was observed in which a group of students successfully transferred and implemented these skills for their undergraduate research project resulting in a UJEMI+ publication (https://jemi.microbiology.ubc.ca/node/188).

**Fig 4.**
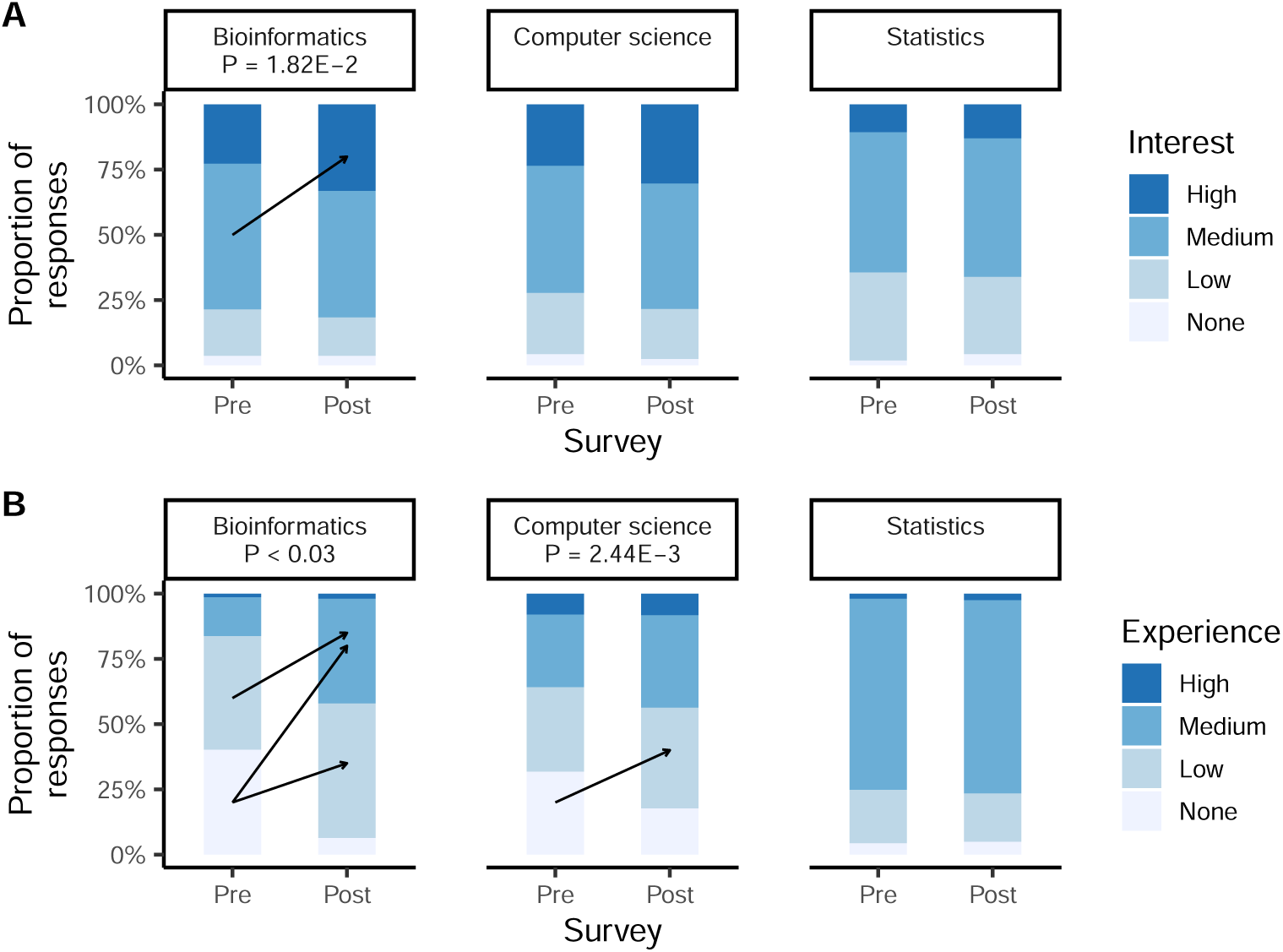
Student interest and experience in data science. Student responses from matched pre- and post-surveys in MICB 301 indicating self-reported interest (N = 143 - 146) and experience (N = 136 -143) in data science areas including bioinformatics and computer science. Numerical responses from 2018/19 were converted to categorical groups from 2017/18 as none (0), low (1-3), medium (4-7), and high (8-10). Arrows and p-values indicate significant response changes by Monte Carlo symmetry tests for paired contingency tables.

Full analysis details are available at https://github.com/EDUCE-UBC/2020_PLOS.

## Conclusion

The EDUCE initiative at UBC was launched in Fall 2017 in response to a clear and present need to expand data science education in the life sciences. The initiative seeks to develop an extensible, progressive, cross-disciplinary and collaborative framework to equip undergraduate students in the life sciences with basic competency in data science. EDUCE learning objectives are achieved through 1) a modular curriculum plugged into existing courses, 2) coordinated co-curricular activities, and 3) a cross-disciplinary community of practice that brings together teaching teams across multiple training levels. The resulting framework allows students to develop progressive data science competency over several years of integrated practice. Initial implementation of EDUCE learning modules and co-curricular activities is focused on ecology and microbiology but can be readily extended to other data sets and discipline-specific software in the life sciences. All EDUCE teaching and learning materials are available on GitHub with the intention of developing an open source community of practice that extends beyond UBC.

While course modules are designed to be extensible, some modification is required prior to implementation in a new course to accommodate individual teaching style, course schedule, and student prior knowledge. This process is made easier through collaboration across the EDUCE community of practice. However, as with any new curriculum, instructor preparation time remains a barrier to implementation of EDUCE modules in some courses. In the future, collaborative platform like GitHub classroom could facilitate collaboration and material exchange between instructors and strengthen the community of practice. Another on-going challenge is student’s prior knowledge. Due to the non-linear nature of MICB degree requirements at UBC, some students come into an intermediate EDUCE courses without prior EDUCE experience while others repeat EDUCE content in multiple courses. To overcome this, supplementary exercises are being developed for students entering EDUCE courses in fourth-year. Although preliminary results indicate that EDUCE is having positive impacts on student interest and learning in data science, further analyses of collected survey data are needed to determine if 1) these perceived changes translate into long-term skill development [20] and if 2) EDUCE curriculum is truly causative of these outcomes.

Despite the need for continued data collection to measure EDUCE impacts, the framework provides a viable approach to data science education in the life sciences that is both flexible and scalable across in person and remote learning platforms. As with any other ability, students need the time and scaffolding to learn data science tools, and the relevant education should not be confined to a single course. The EDUCE community of practice works to transcend siloed educational norms resulting in a zone of proximal development that supports both student and instructor achievement. Given that data science competency also referred to as data literacy is increasingly recognized as a critical but limiting ability in the modern workforce, institutions and granting agencies are encouraged to think about resource allocation beyond the silos of individual courses and departments in order to sustain interdisciplinary teaching and learning frameworks like EDUCE.

## Acknowledgments

The authors wish to thank faculty and instructors for access to courses and support during module integration, including Marcia Graves, Martin Hirst, William Mohn, Sean Crowe, and Jennifer Sibley. We also thank our statistics guru Gaby Cohen Freue, members of the AsDA workshop team including Carolyn Taylor and Biljana Jonoska Stojkova, and teaching assistants for their effort and support throughout the program, including Julia Beni, Jonah Lin, Yue Liu, Nolan Shelley, and David Yin. Special thanks to Zaira Petruf, Kevin Lin, Aria Hahn and Jennifer Bonderoff for getting ECOSCOPE and EDUCE activities organized and off the ground, Sara Harris, Gulnur Birol and Ashley Welsh in the Skylight Science Centre for Learning and Teaching and Jeff Miller in the Centre for Teaching, Learning and Technology for institutional guidance and assessment criteria, and Nancy Heckman in Statistics and Michael Gold and Michael Murphy in MICB for departmental leadership and vision needed to develop and maintain the EDUCE initiative. *Funding:* This work was performed under the auspices of the UBC Teaching and Learning Enhancement Fund (TLEF), the Department of Microbiology and Immunology, and the Natural Science and Engineering Research Council (NSERC) of Canada Collaborative Research and Training Experience (CREATE) ECOSCOPE training program. *Conflict of Interest:* SJH is a co-founder of Koonkie Inc., a bioinformatics consulting company that designs and provides scalable algorithmic and data analytics solutions in the cloud.

