## Supplementary material for "An integrated, modular approach to data science education in the life sciences": Supplmental Text 1

Text S1. Pre- and post-course surveys.

### A. EDUCE pre-course survey - Fall 2017

#### Page 1

##### Introduction

Data science is an increasingly common component of science, technology, engineering, and math (STEM) as well as related careers. Therefore, the U. of British Columbia is employing a number of programs to improve and expand its undergraduate data science curricula. One such program is the Experiential Data Science for Undergraduate Cross-disciplinary Education (EDUCE) initiative, which will impact this course through the introduction of short data science modules.

The goal of this survey is to determine your previous experience with data science before participating in the targeted data science EDUCE modules in this course.

#### For which course are you completing this survey?

- MICB301: Microbial Ecophysiology
- MICB405: Bioinformatics
- MICB421: Experimental Microbiology
- MICB425: Microbial Ecological Genomics

#### Please create an anonymous identifier that is the first three letters of your mother’s first name and the last 4 digits of your cell phone number.

If you do not have a person whom you identify as your mother, please use your father's first name. If you do not have a cell phone, please use your home phone number. For example, if your mother’s name is Nancy and your phone number is 555-555-1828, your identifier would be NAN1828.

Your identifier will be the same throughout this course as well as other courses and workshops, so please follow the setup correctly.

Type here

#### Key terminology

In this section, we would like to determine your previous experience with the terms "data science" and "bioinformatics".

##### Have you heard the term “data science”?

- Yes
- No

##### How would you define data science?

Type here

##### Where have you been exposed to data science? Courses, popular media, books, etc.

Type here

##### Have you heard the term “bioinformatics”?

- Yes
- No

##### How would you define bioinformatics?

Type here

##### Where have you been exposed to bioinformatics? Courses, popular media, books, etc.

Type here

#### Related fields

Please rate your interest and experience in fields related to data science.

#### Please indicate which interfaces and/or programs you have used *prior to this course*.

Check all that apply.

- ☐ Unix command line
- ☐ R
- ☐ mothur
- ☐ QIIME/QIIME2
- ☐ Metagenomic tools (like aligning, assembling, binning)
- ☐ None of the above

#### How would you rate your interest in

| | None | Low | Medium | High |
|---|---|---|---|---|
| computer science? | ○ | ○ | ○ | ○ |
| statistics? | ○ | ○ | ○ | ○ |
| bioinformatics? | ○ | ○ | ○ | ○ |
| microbiology? | ○ | ○ | ○ | ○ |

#### What level of experience do you have in

Levels provide some examples but you may have other experience not listed. Please choose the level you feel fits you best.

| | None | Low (like taking a workshop or doing some readings) | Medium (like taking a course) | High (like taking multiple, high-level courses) | Very high (like publishing in a peer-reviewed journal) |
|---|---|---|---|---|---|
| computer science? | ○ | ○ | ○ | ○ | ○ |
| bioinformatics? | ○ | ○ | ○ | ○ | ○ |
| microbiology? | ○ | ○ | ○ | ○ | ○ |
| statistics? | ○ | ○ | ○ | ○ | ○ |

#### Related coursework

Please detail your previous coursework related to data science.

##### What is your academic major?

Type here

##### Have you previously completed EDUCE module(s)?

- Yes
- No

##### If so, in which course(s)?

Please check all the apply.

- MICB301: Microbial Ecophysiology
- MICB405: Bioinformatics
- MICB421: Experimental Microbiology
- MICB425: Microbial Ecological Genomics

##### Please indicate if you have previously completed, are currently enrolled in, or plan to enroll in any of the following data science-related courses.

###### Microbiology

| | Completed previously | Currently enrolled | Plan to enroll in the future | None |
|---|---|---|---|---|
| MICB301: Microbial Ecophysiology | | | | |
| MICB405: Bioinformatics | | | | |
| MICB421: Experimental Microbiology | | | | |
| MICB425: Microbial Ecological Genomics | | | | |

###### Statistics

| | Completed previously | Currently enrolled | Plan to enroll in the future | None |
|---|---|---|---|---|
| STAT200: Elementary Statistics for Applications | | | | |
| STAT203: Statistical Methods | | | | |
| STAT241: Introductory Probability and Statistics | | | | |
| STAT251: Elementary Statistics | | | | |
| BIOL300: Introduction to Biostatistics | | | | |

###### Computer science

| | Completed previously | Currently enrolled | Plan to enroll in the future | None |
|---|---|---|---|---|
| CPSC100: Computational Thinking | | | | |

| | Completed previously | Currently enrolled | Plan to enroll in the future | None |
|---|---|---|---|---|
| CPSC101: Introduction to Systematic Program Design | ○ | ○ | ○ | ○ |
| CPSC110: Computation, Programs, and Programming | ○ | ○ | ○ | ○ |
| CPSC121: Models of Computation | ○ | ○ | ○ | ○ |
| CPSC301: Computing in the Life Sciences | ○ | ○ | ○ | ○ |

Please list any other relevant courses which you *completed previously*. This could be from the departments listed above (MICB, STAT, CPSC) or any other department with an equivalent course.

Type here

Please list any other relevant courses in which you are *currently enrolled*. This could be from the departments listed above (MICB, STAT, CPSC) or any other department with an equivalent course.

Type here

Please list any other relevant courses in which you *plan to enroll*. This could be from the departments listed above (MICB, STAT, CPSC) or any other department with an equivalent course.

Type here

**Page 6**

#### Related extra-curriculars

Please indicate your experience and interest in other data science-related learning opportunities.

##### Have you participated in any data science-related activities outside of class? Workshops, hackathons, etc.

- Yes
- No

##### If so, please specify.

Type here

##### Are you interested in participating in workshops outside of class to develop or continue to develop your skills in data science?

- Yes
- No

##### Are you aware of any such workshops currently offered at UBC?

- Yes
- No

##### If so, please specify.

Type here

**Page 7**

#### Thank you for taking the time to complete this survey. Please click "Submit" before exiting.

If you have any questions regarding the study or are interested in learning about our findings, please contact Dr. Steven Hallam.

THE UNIVERSITY OF BRITISH COLUMBIA

**About UBC**
Contact UBC
About the University
News
Events
Careers
Make a Gift
Search UBC.ca

**UBC Campuses**
Vancouver Campus
Okanagan Campus

**UBC Sites**
Robson Square
Great Northern Way
Faculty of Medicine Acros
Asia Pacific Regional Offic

### B. EDUCE pre-course survey - 2018-19

**Intro**

**Introduction**

Data science is an increasingly common component of science, technology, engineering, and math (STEM) as well as related careers. Therefore, the U. of British Columbia is employing a number of programs to improve and expand its undergraduate data science curricula. One such program is the Experiential Data Science for Undergraduate Cross-disciplinary Education (EDUCE) initiative, which will impact this course through the introduction of short data science modules.

The goal of this survey is to determine your previous experience with data science before participating in the targeted data science EDUCE modules in this course.

**Course & ID**

For which course are you completing this survey?

- ○ MICB301: Microbial Ecophysiology
- ○ MICB322: Molecular Microbiology Laboratory
- ○ MICB323: Molecular Immunology and Virology Laboratory
- ○ MICB405: Bioinformatics
- ○ MICB421: Experimental Microbiology
- ○ MICB425: Microbial Ecological Genomics
- ○ MICB447: Experimental Molecular Biology
- ○ BIOL436: Integrated Functional Genomics

Please create an anonymous identifier that is the first three letters of your mother’s first name and the last 4 digits of your cell phone number.

> If you do not have a person whom you identify as your mother, please use your father's first name. If you do not have a cell phone, please use your home phone number. For example, if your mother’s name is Nancy and your phone number is 555-555-1828, your identifier would be NAN1828.
>
> Your identifier will be the same throughout this course as well as other courses and workshops, so please follow the setup correctly.

#### Key terms

##### Key terminology

In this section, we would like to determine your previous experience with the terms "data science" and "bioinformatics".

Have you heard the term “data science”?

Yes ○ No ○

How would you define data science?

Where have you been exposed to data science? Courses, popular media, books, etc.

Have you heard the term “bioinformatics”?

Yes ○ No ○

How would you define bioinformatics?

Where have you been exposed to bioinformatics? Courses, popular media, books, etc.

#### Fields

##### Related fields

Please rate your interest and experience in fields related to data science.

Please indicate which interfaces and/or programs you have used *prior to this course*.

- ☐ Unix command line
- ☐ R
- ☐ mothur
- ☐ QIIME/QIIME2
- ☐ Metagenomics tools (like aligning, assembling, binning)
- ☐ None of the above

How would you rate your interest in

| | None | | | Low | | | Medium | | | | High |
|---|---|---|---|---|---|---|---|---|---|---|---|
| | 0 | 1 | 2 | 3 | 4 | 5 | 6 | 7 | 8 | 9 | 10 |
| microbiology? | | | | | | | | | | | |
| bioinformatics? | | | | | | | | | | | |
| statistics? | | | | | | | | | | | |
| computer science? | | | | | | | | | | | |

What level of experience do you have in

| | None | | Low (like taking a workshop or doing some readings) | | | Medium (like taking a course) | | High (like taking multiple, high-level courses) | | | Very high (like publishing in a peer-reviewed journal) |
|---|---|---|---|---|---|---|---|---|---|---|---|
| | 0 | 1 | 2 | 3 | 4 | 5 | 6 | 7 | 8 | 9 | 10 |
| microbiology? | | | | | | | | | | | |
| bioinformatics? | | | | | | | | | | | |
| statistics? | | | | | | | | | | | |
| computer science? | | | | | | | | | | | |

##### Coursework

**Related coursework**

Please detail your previous coursework related to data science.

What is your academic major?

Have you previously completed EDUCE module(s)?

- ○ Yes
- ○ No

If so, in which course(s)?

- ☐ MICB301: Microbial Ecophysiology
- ☐ MICB405: Bioinformatics
- ☐ MICB421: Experimental Microbiology
- ☐ MICB425: Microbial Ecological Genomics

Please indicate if you have previously completed, are currently enrolled in, or plan to enroll in any of the following data science-related courses.

Microbiology

| | Completed previously | Currently enrolled | Plan to enroll in the future | None |
|---|---|---|---|---|
| MICB301: Microbial Ecophysiology | ○ | ○ | ○ | ○ |
| MICB405: Bioinformatics | ○ | ○ | ○ | ○ |
| MICB421: Experimental Microbiology | ○ | ○ | ○ | ○ |
| MICB425: Microbial Ecological Genomics | ○ | ○ | ○ | ○ |
| MICB447: Experimental Molecular Biology | ○ | ○ | ○ | ○ |

Statistics

| | Completed previously | Currently enrolled | Plan to enroll in future | None |
|---|---|---|---|---|

| | Completed previously | Currently enrolled | Plan to enroll in future | None |
|---|---|---|---|---|
| STAT200: Elementary Statistics for Applications | O | O | O | O |
| STAT203: Statistical Methods | O | O | O | O |
| STAT241: Introductory Probability and Statistics | O | O | O | O |
| STAT251: Elementary Statistics | O | O | O | O |
| BIOL300: Introduction to Biostatistics | O | O | O | O |

Computer science

| | Completed previously | Currently enrolled | Plan to enroll in future | None |
|---|---|---|---|---|
| CPSC100: Computational Thinking | O | O | O | O |
| CPSC101: Introduction to Systematic Program Design | O | O | O | O |
| CPSC110: Computation, Programs, and Programming | O | O | O | O |
| CPSC121: Models of Computation | O | O | O | O |
| CPSC301: Computing in the Life Sciences | O | O | O | O |

Please list any other data science relevant courses from the departments listed above (MICB, STAT, CPSC) or any other department with an equivalent course that you

- ☐ Previously completed __________
- ☐ Are currently enrolled in __________
- ☐ Plan to enroll in future __________

**Co-curricular**

**Related co-curriculars**

Please indicate your experience and interest in other data science-related learning opportunities.

Have you participated in any data science-related activities outside of class? Workshops, hackathons, etc.

If so, please specify.

- Yes
- No

Are you interested in participating in workshops outside of class to develop or continue to develop your skills in data science?

- Yes
- No

Are you aware of any such workshops currently offered at UBC?

If so, please specify.

- Yes
- No

Private Policy Terms of Use

Powered by Qualtrics

### C. EDUCE post-course survey - Fall 2017

#### Page 1

##### Introduction

During this semester, you completed one or more data science modules as part of the Experiential Data Science for Undergraduate Cross-disciplinary Education (EDUCE) initiative. The goal of this survey is to measure the impact that these EDUCE modules had on your understanding and experiences in data science.

#### For which course are you completing this survey?

- MICB301: Microbial Ecophysiology
- MICB405: Bioinformatics
- MICB421: Experimental Microbiology
- MICB425: Microbial Ecological Genomics

#### Please create an anonymous identifier that is the first three letters of your mother’s first name and the last 4 digits of your cell phone number.

If you do not have a person whom you identify as your mother, please use your father's first name. If you do not have a cell phone, please use your home phone number. For example, if your mother’s name is Nancy and your phone number is 555-555-1828, your identifier would be NAN1828.

Your identifier will be the same throughout this course as well as other courses and workshops, so please follow the setup correctly.

Type here

#### Key terminology

In this section, we would like to determine how your definitions of "data science" and "bioinformatics" have changed over this semester.

##### Have you heard the term “data science”?

- Yes
- No

##### How would you define data science?

Type here

##### Where have you been exposed to data science? Courses, popular media, books, etc.

Type here

##### Have you heard the term “bioinformatics”?

- Yes
- No

##### How would you define bioinformatics?

Type here

##### Where have you been exposed to bioinformatics? Courses, popular media, books, etc.

Type here

#### Related fields

Please rate your current interest and experience in fields related to data science.

##### How would you rate your interest in

| | None | Low | Medium | High |
|---|---|---|---|---|
| statistics? | ○ | ○ | ○ | ○ |
| microbiology? | ○ | ○ | ○ | ○ |
| computer science? | ○ | ○ | ○ | ○ |
| bioinformatics? | ○ | ○ | ○ | ○ |

##### What level of experience do you have in

| | None | Low (like taking a workshop or doing some readings) | Medium (like taking a course) | High (like taking multiple, high-level courses) | Very high (like publishing in a peer-reviewed journal) |
|---|---|---|---|---|---|
| bioinformatics? | ○ | ○ | ○ | ○ | ○ |
| statistics? | ○ | ○ | ○ | ○ | ○ |
| microbiology? | ○ | ○ | ○ | ○ | ○ |
| computer science? | ○ | ○ | ○ | ○ | ○ |

#### Related coursework

Please indicate how the EDUCE data science module(s) impacted your interest in related coursework.

##### As a result of the EDUCE *data science module(s)* within this course, are you more or less likely to enroll in the following courses?

##### Microbiology

| | Less | More | No change | N/A, previously completed course |
|---|---|---|---|---|
| MICB301: Microbial Ecophysiology | ○ | ○ | ○ | ○ |
| MICB405: Bioinformatics | ○ | ○ | ○ | ○ |
| MICB421: Experimental Microbiology | ○ | ○ | ○ | ○ |
| MICB425: Microbial Ecological Genomics | ○ | ○ | ○ | ○ |

##### Statistics

| | Less | More | No change | N/A, previously completed course |
|---|---|---|---|---|
| STAT200: Elementary Statistics for Applications | ○ | ○ | ○ | ○ |
| STAT203: Statistical Methods | ○ | ○ | ○ | ○ |
| STAT241: Introductory Probability and Statistics | ○ | ○ | ○ | ○ |
| STAT251: Elementary Statistics | ○ | ○ | ○ | ○ |
| BIOL300: Introduction to Biostatistics | ○ | ○ | ○ | ○ |

##### Computer science

| | Less | More | No change | N/A, previously completed course |
|---|---|---|---|---|
| CPSC100: Computational Thinking | ○ | ○ | ○ | ○ |
| CPSC101: Introduction to Systematic Program Design | ○ | ○ | ○ | ○ |
| CPSC110: Computation, Programs, and Programming | ○ | ○ | ○ | ○ |
| CPSC121: Models of Computation | ○ | ○ | ○ | ○ |
| CPSC301: Computing in the Life Sciences | ○ | ○ | ○ | ○ |

##### Please list any other relevant courses in which you are more or less likely to enroll based on your experiences in the *data science modules* in this course.

Type here

#### Related extra-curriculars

Please indicate your current experience and interest in other data science-related learning opportunities.

##### Have you taken a data science workshop outside of class *this semester*?

- Yes
- No

##### If so, please specify.

Type here

##### Are you interested in participating in workshops outside of class to develop or continue to develop your skills in data science?

- Yes
- No

##### If so, which one(s) interest you?

Please check all that apply.

##### Introductory

- Introduction to R
- Data carpentry: Basic data organization, manipulation, and visualization in R
- Software carpentry: Programming in Python, the Unix shell, and version control with Git

##### Intermediate

- Working with tabular data in R
- Graphics with ggplot in R
- Statistical models in R
- Functions, workflows, and reproducible research in R and Git

##### Microbiome

- Amplicon sequence data analysis in mothur or QIIME
- Microbiome analysis in R
- Metagenomics analysis
- Metatranscriptomics analysis

#### Demographics

This section is entirely optional and for research purposes only.

##### Do you identify as a racial or ethnic minority?

- Yes
- No
- Prefer not to answer

##### Do you identify as a first-generation university student?

*i.e.* Neither of your parent(s) or guardian(s) attended college.

- Yes
- No
- Prefer not to answer

##### Do you identify as a non-traditional university student?

This includes those who:

- Delay enrollment after high school by more than a year or
- Attend part-time for at least part of the academic year or
- Work full-time while enrolled or
- Have dependents other than a spouse

- Yes
- No
- Prefer not to answer

#### Page 8

##### Thank you for taking the time to complete this survey.

Please click "Submit" and you will be redirected to a page asking if you are willing to participate in a follow-up survey in the future.

If you have any questions regarding the study or are interested in learning about our findings, please contact Dr. Steven Hallam.

THE UNIVERSITY OF BRITISH COLUMBIA

**About UBC**
Contact UBC
About the University
News
Events
Careers
Make a Gift
Search UBC.ca

**UBC Campuses**
Vancouver Campus
Okanagan Campus

**UBC Sites**
Robson Square
Great Northern Way
Faculty of Medicine Acros
Asia Pacific Regional Offic

### D. EDUCE post-course survey - 2018-19

**Intro**

**Introduction**

During this semester, you completed one or more data science modules as part of the Experiential Data Science for Undergraduate Cross-disciplinary Education (EDUCE) initiative. The goal of this survey is to measure the impact that these EDUCE modules had on your understanding and experiences in data science.

**Course & ID**

For which course are you completing this survey?

- ○ MICB301: Microbial Ecophysiology
- ○ MICB322: Molecular Microbiology Laboratory
- ○ MICB323: Molecular Immunology and Virology Laboratory
- ○ MICB405: Bioinformatics
- ○ MICB421: Experimental Microbiology
- ○ MICB425: Microbial Ecological Genomics
- ○ MICB447: Experimental Molecular Biology
- ○ BIOL436: Integrated Functional Genomics

Please create an anonymous identifier that is the first three letters of your mother’s first name and the last 4 digits of your cell phone number.

> If you do not have a person whom you identify as your mother, please use your father's first name. If you do not have a cell phone, please use your home phone number. For example, if your mother’s name is Nancy and your phone number is 555-555-1828, your identifier would be NAN1828.
>
> Your identifier will be the same throughout this course as well as other courses and workshops, so please follow the setup correctly.

[ ]

**Key terms**

**Key terminology**

In this section, we would like to determine how your definitions of "data science" and "bioinformatics" have changed over this semester.

Have you heard the term “data science”?

Yes ○ No ○

How would you define data science?

Where have you been exposed to data science? Courses, popular media, books, etc.

Have you heard the term “bioinformatics”?

Yes ○ No ○

How would you define bioinformatics?

Where have you been exposed to bioinformatics? Courses, popular media, books, etc.

**Related fields**

**Related fields**

Please rate your current interest and experience in fields related to data science.

How would you rate your interest in

| | None | | | Low | | | Medium | | | | High |
|---|---|---|---|---|---|---|---|---|---|---|---|
| | 0 | 1 | 2 | 3 | 4 | 5 | 6 | 7 | 8 | 9 | 10 |
| microbiology? | | | | | | | | | | | |
| bioinformatics? | | | | | | | | | | | |
| statistics? | | | | | | | | | | | |
| computer science? | | | | | | | | | | | |

What level of experience do you have in

| | None | | Low (like taking a workshop or doing some readings) | | | Medium (like taking a course) | | High (like taking multiple, high-level courses) | | | Very high (like publishing in a peer-reviewed journal) |
|---|---|---|---|---|---|---|---|---|---|---|---|
| | 0 | 1 | 2 | 3 | 4 | 5 | 6 | 7 | 8 | 9 | 10 |
| microbiology? | | | | | | | | | | | |
| bioinformatics? | | | | | | | | | | | |
| statistics? | | | | | | | | | | | |
| computer science? | | | | | | | | | | | |

**Related coursework**

**Related coursework**

Please indicate how the EDUCE data science module(s) impacted your interest in related coursework.

As a result of the EDUCE *data science module(s)* within this course, are you more or less likely to enroll in the following courses?

Microbiology

| | Less | More | No change | N/A, previously completed course |
|---|---|---|---|---|
| MICB301: Microbial Ecophysiology | O | O | O | O |
| MICB405: Bioinformatics | O | O | O | O |

| | Less | More | No change | N/A, previously completed course |
|---|---|---|---|---|
| MICB421: Experimental Microbiology | O | O | O | O |
| MICB425: Microbial Ecological Genomics | O | O | O | O |
| MICB447: Experimental Molecular Biology | O | O | O | O |

Statistics

| | Less | More | No change | N/A, previously completed course |
|---|---|---|---|---|
| STAT200: Elementary Statistics for Applications | O | O | O | O |
| STAT203: Statistical Methods | O | O | O | O |
| STAT241: Introductory Probability and Statistics | O | O | O | O |
| STAT251: Elementary Statistics | O | O | O | O |
| BIOL300: Introduction to Biostatistics | O | O | O | O |

Computer science

| | Less | More | No change | N/A, previously completed course |
|---|---|---|---|---|
| CPSC100: Computational Thinking | O | O | O | O |
| CPSC101: Introduction to Systematic Program Design | O | O | O | O |
| CPSC110: Computation, Programs, and Programming | O | O | O | O |
| CPSC121: Models of Computation | O | O | O | O |
| CPSC301: Computing in the Life Sciences | O | O | O | O |

Please list any other relevant courses in which you are more or less likely to enroll based on your experiences in the *data science modules* in this course.

**Co-curricular**

**Related co-curriculars**

Please indicate your current experience and interest in other data science-related learning opportunities.

Have you taken a data science workshop outside of class *this semester*?

If so, please specify.

- ○ Yes [ ]
- ○ No

Are you interested in participating in workshops outside of class to develop or continue to develop your skills in data science?

- ○ Yes
- ○ No

If so, which one(s) interest you? Please check all that apply.

Introductory

- ☐ Introduction to R: RStudio base R graphics and data
- ☐ Advanced introduction to R: RStudio graphics with ggplot and data manipulation with dplyr
- ☐ R for researchers: RStudio reproducible research with GitHub and data manipulation with dplyr

Intermediate

- ☐ Working with data: RStudio complex data manipulation with dplyr and tidyr
- ☐ Graphics with ggplot: RStudio customizable graphics with ggplot
- ☐ Statistical model: RStudio model assumptions, outputs, and functions
- ☐ Reproducible research: RStudio and GitHub integration

Advanced

- ☐ Advanced R programming: RStudio function, class, and package creation
- ☐ Microbiome data analysis in mothur: from fastq to OTU table
- ☐ Microbiome data analysis in QIIME2: from fastq to ASV table
- ☐ Microbiome data analysis in R: uses of vegan and phyloseq in microbiome research

**Demographics**

This section is entirely optional and for research purposes only.

Do you identify as a racial or ethnic minority?

- Yes
- No
- Prefer not to answer

Do you identify as a first-generation university student?

*i.e.* Neither of your parent(s) or guardian(s) attended college.

- Yes
- No
- Prefer not to answer

Do you identify as a non-traditional university student?

This includes those who:

Delay enrollment after high school by more than a year or
Attend part-time for at least part of the academic year or
Work full-time while enrolled or
Have dependents other than a spouse

- Yes
- No
- Prefer not the answer

Private Policy Terms of Use

Powered by Qualtrics

Text S1. Pre- and post-course surveys.
